# Estimation of Mediation Effect for High-dimensional Omics Mediators with Application to the Framingham Heart Study

**DOI:** 10.1101/774877

**Authors:** Tianzhong Yang, Jingbo Niu, Han Chen, Peng Wei

## Abstract

Environmental exposures can regulate intermediate molecular phenotypes, such as gene expression, by different mechanisms and thereby lead to various health outcomes. It is of significant scientific interest to unravel the role of potentially high-dimensional intermediate phenotypes in the relationship between environmental exposure and traits. Mediation analysis is an important tool for investigating such relationships. However, it has mainly focused on low-dimensional settings, and there is a lack of a good measure of the total mediation effect. Here, we extend an R-squared (Rsq) effect size measure, originally proposed in the single-mediator setting, to the moderate- and high-dimensional mediator settings in the mixed model framework. Based on extensive simulations, we compare our measure and estimation procedure with several frequently used mediation measures, including product, proportion, and ratio measures. Our Rsq measure has small bias and variance under the correctly specified model. To mitigate potential bias induced by non-mediators, we examine two variable selection procedures, i.e., iterative sure independence screening and false discovery rate control, to exclude the non-mediators. We evaluate the consistency of the proposed estimation procedures and introduce a resampling-based confidence interval. By applying the proposed estimation procedure, we find that more than half of the aging-related variations in systolic blood pressure can be explained by gene expression profiles in the Framingham Heart Study.

## 1 Introduction

Understanding the relationships between an environmental risk factor and different health traits through molecular phenotypes, such as gene expression (GE) and DNA methylation, can provide mechanistic insights into disease etiology and exposure biology. Specifically, an environmental risk factor may lead to epigenetic changes, such as changes in DNA methylation, which then alter DNA accessibility and chromatin structure, and thereby regulate GE and further downstream molecular phenotypes pertinent to the disease process. Modern epidemiological studies are capable of measuring a large number of markers, from tens of thousands of GEs to nearly a million CpG sites in DNA methylation studies. There is growing evidence that many of these intermediate phenotypes could lie in the pathway between environmental exposure and downstream health outcomes (Ladd-Acosta and Fallin, 2016). It is thus of great scientific interest to quantify the total amount of variation in health traits that is explained by environmental risk factors through the intermediate phenotypes. Mediation analysis is a natural approach to explore such relationships, which can help researchers delineate why and how two variables (dependent variable and independent variable) are related (MacKinnon, 2008).

Our motivating scientific question here is how aging affects different health traits through molecular phenotypes. Specifically, we are interested in exploring the mediating role of GEs in the pathway between aging and two health traits, blood pressure (BP) and lung function. As an important risk factor for a wide range of health conditions, aging can be regarded as a proxy of lifestyle, oxidative stress, or other accumulated environmental risk factors. Researchers have found that GE profiles are associated with the aging process in various biological pathways, notably those involving overexpression of inflammation and immune response genes and underexpression of collagen and energy metabolism genes (De Magalhaes et al., 2009; Weindruch et al., 2002). On the other hand, a decrease in lung function and increase in systolic BP were found to be associated with many age-related changes, including inflammation and altered immunity, and these changes may be reflected on the molecular level (Torre-Amione, 2005; Lowery et al., 2013; Huan et al., 2015; Obeidat et al., 2015). Instead of exploring the mediating effect of a particular gene, we intend to quantify the overall role of potentially high-dimensional GEs in mediating the relationship between aging and health traits, i.e., the total mediation effect. As far as we know, the existing total mediation effect size measures have been studied under low-dimensional settings and all of them are based on the difference in means, i.e., first moment estimand (to be detailed later). Less attention has been given to the moderate- and high-dimensional settings (Miočević et al., 2018; Preacher and Kelley, 2011), although such a measure may be especially useful in guiding further more specific pathway analyses and providing mechanistic insights.

To fill in the gap, we extend a total mediation effect size measure, the R-squared effect size measure (Rsq measure), which was originally proposed in a single-mediator model by Fairchild et al. (2009), to the multiple- and high-dimensional mediator models. Briefly, the Rsq measure is the amount of variance of the dependent variable that is common to both the independent variable and the mediator(s), derived from commonality analysis (Seibold and McPhee, 1979; Ray-Mukherjee et al., 2014). As an estimand based on variation, the Rsq measure provides an alternative to existing measures, especially in the presence of possible opposite directions of mediation effects in the literature (Huang et al., 2018) and the motivating example (Fig S3). To further deal with high-dimensional candidate mediators, we establish a consistent estimation procedure of the Rsq measure. Briefly, we use a variable selection method with the oracle property to filter out the non-mediators that bias the Rsq measure, and then we adopt a mixed-effect model to obtain stable Rsq estimates. The variable selection step in the high-dimensional mediation analysis is not trivial because the identity of true mediators and biological mechanisms in real data are rarely known *a priori*. Therefore, in addition to theoretical justification, we conduct extensive simulations to evaluate this estimation procedure from various perspectives, including bias, variance, finite sample performance of consistency, and the coverage probability of the confidence interval (CI). We show that our method has an all-around performance.

We apply the proposed estimation procedure to answer our motivating question using the Framingham Heart Study (FHS) data, which contains a total of 17,873 candidate genes with corresponding GEs, 1711 subjects for BP evaluation, and 1378 subjects for lung function evaluation. Since the GE levels in the FHS were measured at the same time, we assume undirected correlations among the GE levels, following Huang and Pan (2016) and Boca et al. (2013). Nonetheless, we demonstrate that the Rsq measure is also viable to use with directed correlation among mediators. The main consideration of our study is the magnitude of the total mediation effect, instead of hypothesis testing that considers whether the effect is present or not. In addition, we do not pursue causal interpretations as the assumptions are often violated and difficult to verify in the real-data application (VanderWeele, 2016). The rest of the article is organized as follows. Section 2 reviews mediation analysis, introduces the new Rsq effect size measure and proposes a mixed-model based estimation procedure, coupled with variable selection in high-dimensional settings. Section 3 presents extensive simulations, followed by application to the FHS dataset in Section 4. We end with discussion and future directions in Section 5.

## 2 Methods

### 2.1 Review of mediation analysis and three effect size measures

A mediation model (Fig. 1) consists of the following equations. Without loss of generality, we assume the dependent variable and mediators are centered.

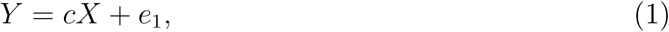

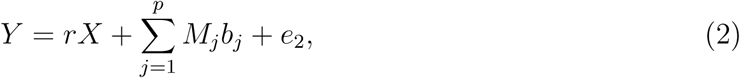

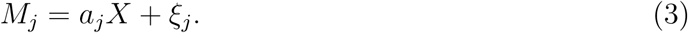

*p* is the total number of mediators. When *p* = 1, it corresponds to a single-mediator model (Fig.1.A); otherwise, it corresponds to a multiple-mediator model (Fig.1.B). *Y* is the continuous dependent variable; *X* is the independent variable; *M*_*j*_ is the *j*^*th*^ mediator; *e*_1_, *e*_2_, and *ξ*_*j*_ are residuals for each equation; *a*_*j*_, *b*_*j*_, *r* and *c* are regression coefficients, usually estimated by the maximum likelihood estimation (MLE) method. Parameter *c* is the total effect and *r* is the direct effect.

**Figure 1.**
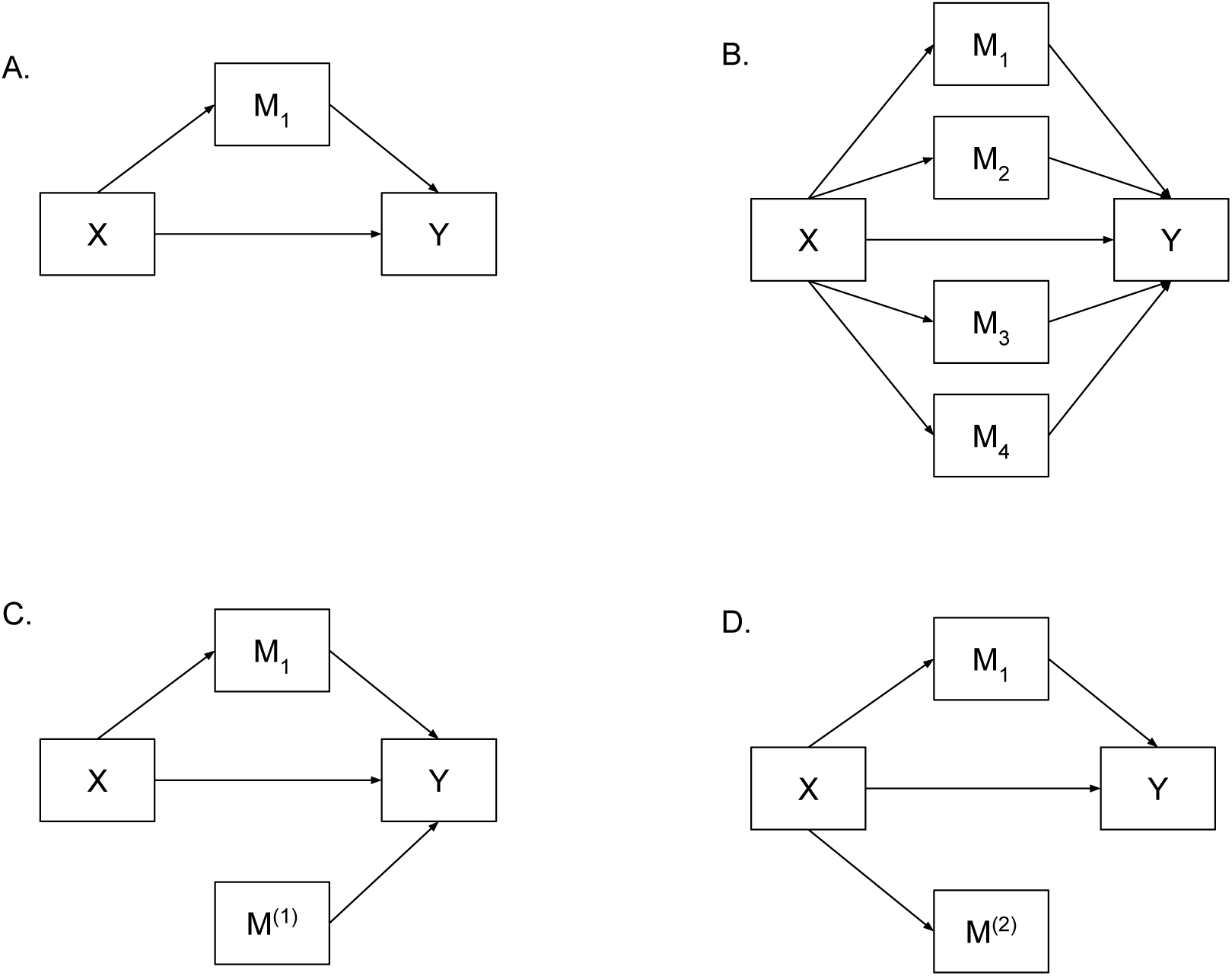
Demonstration of mediation analysis. X is the independent variable, Y is the dependent variable, and *M*_*j*_ is the true mediator; A. Single-mediator model; B. Multiple-mediator model; C shows *M* ^(1)^ that is not associated with X, but with Y; D demonstrates *M* ^(2)^ that is associated with X, but not associated with Y after adjusting for X.

Product, proportion, and ratio measures, all based on the difference in means, are among the most commonly seen total mediation effect measures in the literature. The product measure is 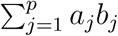. It is also the natural indirect effect under the potential outcome framework with strong causal inference and model assumptions (Imai and Yamamoto, 2013; VanderWeele and Vansteelandt, 2014). The proportion mediated is defined as the proportion of total effect mediated by 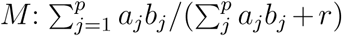; the ratio mediated is 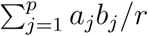. All three measures are sensitive to the direction of effects through different individual mediation pathways. In an extreme example, *a*_*j*_*b*_*j*_ from individual pathways have different directions and thus cancel out, leading to an implication that there is no mediation effect at all. Additionally, both the proportion and ratio measures require a sample size larger than 500 to obtain stable estimates even under low-dimensional settings (MacKinnon, 2008).

### 2.2 Rsq measure under a single-mediator model

Compared with the aforementioned total mediation effect size measures, the Rsq measure has not drawn much attention and has been only formally developed under the single-mediator model. It is defined as the proportion of variance in dependent variable Y explained by the independent variable X through the mediators (Fairchild et al., 2009), as demonstrated via a Venn diagram in Fig. S1. In addition to model (1) and (2), model (4) with *p* = 1 is required for calculation of the Rsq measure:

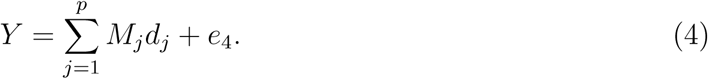

*d*_*j*_ is the regression coefficient and *e*_4_ is the residual. The Rsq measure is defined as 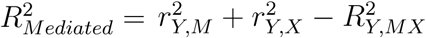, where *r*_*Y,M*_ = *cor*(*Y, M*) is the proportion of variance in Y explained by M in model (4), *r*_*Y,X*_ = *cor*(*Y, X*) is the proportion of variance in Y explained by X in model (1), and 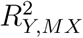 is the proportion of variance in Y explained by M and X in model (2). The three components can be estimated by the MLE using fixed-effect models, i.e., treating all the coefficients as fixed. Different from the above first-moment measures, the Rsq is a difference-in-variance measure. It has been recognized to have many characteristics of a good measure of effect size. For example, the Rsq measure has a stable performance for sample sizes *>* 100 (Fairchild et al., 2009), it increases as the mediation effect approaches total effect, and it is possible to construct a CI estimate (Preacher and Kelley, 2011).

### 2.3 Extension: Rsq measure under the multiple-mediator model

We extend the Rsq measure to the multiple-mediator model, defined as:

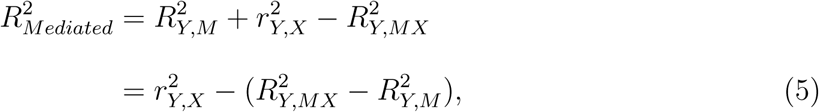

where *r*_*Y,X*_ = *cor*(*Y,X*), 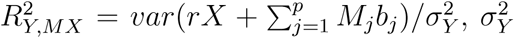 is the variance of Y, and 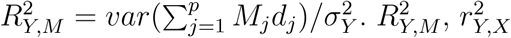, and 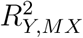 have the same meaning as in the single mediator models and the corresponding models are (4), (1) and (2) with *p >* 1. 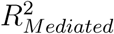 can be interpreted as that in commonality analysis (Seibold and McPhee, 1979): the variance that is common to both the independent variable and the mediator(s), which is evaluated by the difference in the variance of the dependent variable that is explained by the exposure 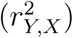 and the additional variance that can be explained by the exposure after taking into account the mediators 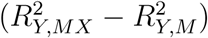, i.e., represented by equation (5). A major concern of using the Rsq measure under a single-mediator model is that it can be negative (Preacher and Kelley, 2011). However, we show that it is probably overstated for both single- and multiple-mediator models, with more discussion in Supplementary Materials Section 1.2. Moreover, we have established several additional appealing properties for the Rsq measure, including (1) invariance to certain transformations, such as principal component analysis (Supplementary Materials Section 1.2.4 Proposition 6), (2) adaptability to a complex casual structure (Supplementary Materials Section 1.3), and (3) no induced bias even if certain types of non-mediators are included (Supplementary Materials Section 1.2.4 Proposition 4). Another closely related measure is the shared over simple effect (SOS) (Lindenberger and Potter, 1998) measure, which is defined as 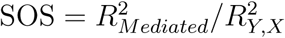. SOS is a relative measure of 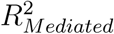. It is the proportion of exposure-related variance in the outcome that is shared with the mediator. The relationships among the Rsq, SOS, product, proportion, and ratio measures are described in Supplementary Materials Section 1.2.2. Interestingly, we find that SOS is closely related to the proportion measure, although they have different interpretations: SOS monotonically increases with the proportion mediated; on the other hand, it is able to capture some bi-directional mediation effects when the proportion measure cannot.

#### 2.3.1 Modelling and estimation

In order to obtain stable estimation under high-dimensional settings, we use mixed-effect model for improved statistical efficiency, as shown later in the numerical examples. Specifically, we assume that the coefficients for the mediators in models (2) and (4) are random effects. In model (2), *b*_*j*_ is assumed to follow a normal distribution *b*_*j*_ ∼ *N*(0, *τ*_1_) for *j* = 1, 2, …, *p* and *e*_2_ ∼ *N*(0, *ϕ*_1_), thus

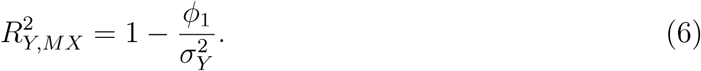

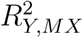 can be interpreted as one minus the variance that is unexplained by the independent variable and mediators. Similarly, in model (4), we assume *d*_*j*_ ∼ *N*(0, *τ*_2_) for *j* = 1, 2, …, *p* and *e*_4_ ∼ *N*(0, *ϕ*_2_), such that

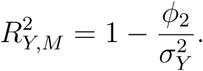

We estimate *τ*_1_,*τ*_2_, *ϕ*_1_ and *ϕ*_2_ by the restricted maximum likelihood method, which is consistent under mild conditions (Cressie and Lahiri, 1993). We estimate 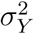 by a consistent estimator 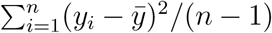, where 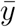 is the sample mean. Note that we avoid the direct use of the estimation of a total of 2*p* coefficients (*β*_1_, …, *β*_*p*_, *d*_1_, …, *d*_*p*_); instead, we use two parameters (*ϕ*_1_ and *ϕ*_2_) to calculate 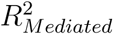. The estimation of latter is robust to the misspecification of the distribution of the random effects; it has been supported by multiple theoretical studies and real-data analysis (Verbeke and Lesaffre, 1997; McCulloch and Neuhaus, 2011; Yang et al., 2019). Finally, 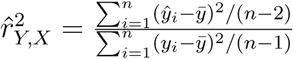 where *ŷ*_*i*_ is the fitted value estimated by MLE in model (1).

When *p << n*, it is also feasible to estimate the three Rsq components by MLE in the fixed-effect models, and we evaluate its performance in the simulation study for comparison.

#### 2.3.2 Mediator variable selection

In the traditional mediation analysis, the mediating variables are hypothesized and selected based on specific research questions and subject matter knowledge. However, hypothesizing and identifying the true mediators becomes much harder in the high-dimensional settings where the bias for estimating the total mediation effects can be induced by failing to identify the true mediators. Inspired by real-data applications, we differentiated the problem into three categories. The first category is the scenario in which the variables falsely identified as mediators are not associated with the exposure, and thus, not in the pathway from the exposure to the outcome (Fig 1.C). For example, some genes influencing lung function are not in the pathway between aging and lung function but others, such as a pathway between smoking and lung function. We denote the set of such variables as **M**^(**1**)^ = {*M*_*j*_: *b*_*j*_ ≠ 0, *a*_*j*_ = 0}. Supplementary Materials Section 1.2.3, Proposition 4, shows that 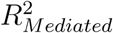 is not biased when **M**^(**1**)^ are included. The second category is the scenario in which the variables are associated with the exposure, but not the outcome after adjusting for the exposure (Fig 1.D). For example, collagen synthesis is age-related, but genes associated with collagen synthesis may not influence BP. We denote the set of such variables as **M**^(**2**)^ = {*M*_*j*_: *a*_*j*_ ≠0, *b*_*j*_ = 0}. We show that the 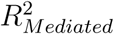 estimate is biased when **M**^(**2**)^ are included as mediators in Supplementary Materials Section 1.2.3, Proposition 5, as well as the simulation study. Mathematically, the bias comes from 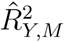, where part of the variance of X is falsely added due to the inclusion of **M**^(**2**)^. The third category is the scenario in which noise variables (*b* = 0 and *a* = 0) are included, for example, genes not associated with aging or the two health traits. The inclusion of noise variables does not influence the estimation of 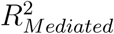 in terms of bias because of the same reason as **M**^(**1**)^. In the steps recommended by Baron and Kenny (1986) for mediation analysis, **M**^(**1**)^, **M**^(**2**)^, and noise variables are not considered as mediators, and thus should be excluded from mediation analysis. One promising feature of our Rsq measure under high-dimensional settings is its robustness to the inclusion of **M**^(**1**)^ and noise variables. However, the inclusion of **M**^(**2**)^ is problematic, which we use a variable selection method to filter out in model (2). For illustration purposes, we denote the true mediators as **M**, the putative mediating variables in the initial assessment as **M**_0_, and the variables included in the final mediation model as 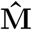 in the following context.

##### Sure independence screening (SIS)

To make the high-dimensional problem solvable, we assume that the true mediators are sparse in our motivating question. We adopt iterative SIS, an extension of SIS, to conduct mediator variable selection. Fan and Lv (2008) introduced SIS in the context of ultrahigh-dimensional linear models, which has a sure screening property, i.e., with probability tending to 1, the independence screening technique retains all of the important predictors in the model under certain conditions. The SIS was used in high-dimensional mediation analysis with a focus on hypothesis testing by Zhang et al. (2016). For our purposes, we use iterative SIS to handle cases where the regularity conditions of SIS fail due to the existence of **M**^(2)^. For example, some genes maybe jointly uncorrelated with the health traits, but have higher marginal correlations with those traits than true mediators. We establish the consistency of the mixed-model approach to 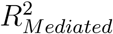 estimation coupled with iterative SIS-minimax concave penalty (MCP) in Supplementary Materials Section 1.2.5, i.e., as *n* → *∞*, 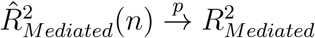

##### Controlling FDR in testing the marginal association

Another common practice for filtering out the undesirable variables is to test the marginal association of each potential mediator with *Y* based on the false discovery rate (FDR) control (Efron, 2012). Boca et al. (2013) discussed how to test different nulls for mediation analysis by controlling the family-wise error rate when hundreds or thousands of mediators are involved. Controlling the FDR is known to be more powerful than controlling the family-wise error rate. Assuming that the true mediators are marginally associated with *Y*, we calculate the FDR-adjusted p-values for the *a*_*j*_’s and *b*_*j*_’s in model (3) and the marginal model between independent variable and mediators (*Y* ∼ *b*_*j*_*M*_*j*_), respectively. If either FDR is larger than 0.1, the variable is excluded from the analysis.

#### 2.3.3 Bootstrap-based confidence interval

To obtain valid post-selection inference, we split the data into two halves, using one half to select the true mediator(s) and the other half to estimate 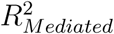(Sun and Bull, 2005; Lee et al., 2016). We use the nonparametric bootstrap method to calculate the percentile CI with details provided in Supplementary Materials Section 1.4.

## 3 Simulation Study

We conducted extensive simulations to assess the performance of the proposed 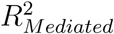 measure, along with the product, proportion, and ratio measures. We simulated data using the same set of coefficients across 500 replications to assess bias, variance, and finite-sample performance of consistency (Supplementary Materials Section 1.5.4), such that the true values of 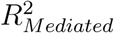 under the mixed-effect models are equal to our derived functions of parameters (Equation (S4)). We used our proposed estimation method, the mixed-effect model, for 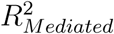 in the following simulation settings. We lightly touched on using the fixed-effect model to estimate the 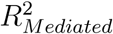, and we only presented its performance in Simulation setting I, low-dimensional setting. In addition to the iterative SIS and FDR control, we examined the Lasso regression for simultaneous variable selection and estimation. Lastly, we evaluated the coverage probability of the proposed bootstrap procedure for CI (Supplementary Materials Section 1.5.5).

### 3.1 Simulation settings

#### 3.1.1 Simulation setting I: bias and variance

We evaluated the bias and variance of different types of total mediation effect measures under both low- and high-dimensional settings. We are interested in the performance of our proposed Rsq measure 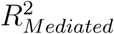 when mediation effects are in the same or different directions and when three types of previously defined non-mediators are included. In addition, we evaluated its performance when a directed causal structure among mediators existed in the low-dimensional setting and when the random effects followed a non-Gaussian distribution under the high-dimensional setting. The simulation set-ups and results for the low-dimensional settings are included in Supplementary Materials Section 1.4. For high-dimensional settings, data were generated using model (2) and (3). The sample size was *n* = 1500, *p* = 1500, *e*_2_ ∼ *N*(0, 1), *X* ∼ *N*(0, 1), and *r* = 1. There were *p* variables in **M**_**0**_, and *ξ* = (*ξ*_1_, *ξ*_2_, …, *ξ*_*p*_) ∼ *N*(0, **D**_*p×p*_), where **D**_*p×p*_ is the identity matrix. The coefficients were estimated by the mixed model or the Lasso regression. The following ten scenarios were assessed: the true values of each measure are provided in Table S2 and S3, where (H1) and (H5) represent the scenarios of correct model specification, (H2), (H3), and (H4) represent the scenarios of including the three types of non-mediators.

(H1) All variables included were true mediators 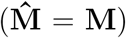 with different directions: *a*_*j*_ ∼ *N*(0, 0.2), *b*_*j*_ ∼ *N*(0, 0.2) for *j* = 1, …, 150;

(H2) Adding additional 1350 **M**^(2)^ to (H1): *a*_*j*_ ∼ *N*(0, 0.2), *b*_*j*_ = 0 for *j* = 151, …, 1500;

(H3) Adding additional 1350 **M**^(1)^ to (H1): *a*_*j*_ = 0, *b*_*j*_ ∼ *N*(0, 0.2) for *j* = 151, …, 1500;

(H4) Adding additional 1350 noise variables to (H1): *a*_*j*_ = 0, *b*_*j*_ = 0 for *j* = 151, …, 1500;

(H5) All variables included were mediators with positive directions: *a*_*j*_ and *b*_*j*_ were the absolute values of the coefficients in (H1);

(H6) - (H10) Same as (H1) to (H5), except that *a*_*j*_’s and *b*_*j*_’s followed a scaled t-distribution with the degree of freedom equal to 1.

#### 3.1.2 Simulation setting II: variable selection

The existence of **M**^(2)^ could bias the estimation of our proposed measure, thus we evaluated two commonly used variable selection methods (iterative SIS and marginal association tests controlling FDR) by examining their impact on the bias, standard deviation (SD), and mean square of error (MSE) of the estimation of 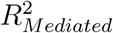. We set *n* = 1500, *r* = 3, *e*_2_ ∼ *N*(0, 1), and *X* ∼ *N*(0, 1); the initial size of **M**_**0**_ is *p* = 1500; **D**_*p×p*_ is the identity matrix. We evaluated the variable selection performance by using (V1) and (V2), representing the scenarios of including two types of non-mediators **M**^(1)^ and **M**^(2)^:

(V1) There were *p*_1_ true mediators, and the rest were **M**^(2)^: *a*_*j*_ ∼ *N*(0, 0.2) for *j* = 1, …, 1500, and *b*_*j*_ ∼ *N*(0, 0.2) for *j* = 1, …, *p*_1_, *b*_*j*_ = 0 for *j* = *p*_1_ + 1, …, 1500;

(V2) There were *p*_1_ true mediators, and the rest were **M**^(1)^: *b*_*j*_ ∼ *N*(0, 0.2) for *j* = 1, …, 1500, and *a*_*j*_ ∼ *N*(0, 0.2) for *j* = 1, …, *p*_1_, *a*_*j*_ = 0 for *j* = *p*_1_ + 1, …, 1500.

We varied *p*_1_ at 0, 15, 75, 150, and 300, corresponding to 0, 1, 5, 10, and 20 percent of the true mediators, respectively. The variable selection was performed in the first half of the data, and the estimation of 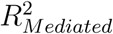 was in the second half, which we benchmarked against the 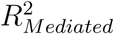 estimate without variable selection 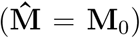 and the Lasso regression product estimate based on all data.

### 3.2 Simulation results

#### 3.2.1 Simulation setting I: bias and variance assessment

Table 1 presents the bias and variance under the high-dimensional settings, i.e., (H1) to (H5). When the model consisted of the true mediators (H1, H5), non-mediators **M**^(1)^ (H3), and noise variables (H4), the 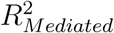 estimators had very small bias and variance. Estimators of the product, proportion, and ratio mediated had relatively high bias when *n* = *p* under scenarios (H2) to (H4), probably because it required estimating a large number of coefficients. In addition, the 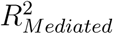 estimators were biased under scenario (H2) as expected, suggesting the importance of excluding non-mediators **M**^(2)^. We further confirmed that our normal assumption on the distribution of random effects was quite robust to misspecification (Table S3). The simulation results for the low-dimensional settings are presented and discussed in Supplementary Materials Section 1.5.1, where we found that mixed-effect models had a slightly better performance in estimating 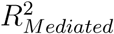 and SOS, compared with fixed-effect models; however, fixed-effect models had a better performance in estimating the product, proportion, and ratio mediated (Table S1).

**Table 1.** Bias and standard deviation under high-dimensional settings (Simulation setting I): bias in the first row, and standard deviation in the second row for each scenario. ab: product measure; prop: proportion measure. (Lasso) indicates that the estimation is based on the Lasso regression; otherwise, it is estimated by a mixed-effect model. The true values are presented in Supplementary Materials Table S2.

| | $R^2_{Mediated}$ | SOS | ab | ab (Lasso) | prop | prop (Lasso) | ratio | ratio (Lasso) |
| --- | --- | --- | --- | --- | --- | --- | --- | --- |
| H1 | 0.0006<br>(0.0181) | 0.0013<br>(0.0370) | -0.0084<br>(0.2846) | -0.0324<br>(0.2744) | 0.0001<br>(0.0833) | 0.0069<br>(0.0795) | -0.0107<br>(0.1161) | -0.0231<br>(0.1117) |
| H2 | 0.0146<br>(0.0184) | 0.0299<br>(0.0375) | 0.1602<br>(0.6463) | -0.0359<br>(0.2604) | -0.0493<br>(0.1960) | 0.0058<br>(0.0777) | 0.0075<br>(0.2886) | -0.0212<br>(0.1165) |
| H3 | 0.0006<br>(0.0071) | 0.0053<br>(0.0653) | 0.0923<br>(0.7443) | 0.0547<br>(0.7547) | -0.0552<br>(0.2520) | -0.0520<br>(0.2495) | -0.0013<br>(0.2983) | -0.0315<br>(0.3392) |
| H4 | 0.0047<br>(0.0198) | 0.0095<br>(0.0403) | 0.1421<br>(0.2613) | -0.0347<br>(0.2519) | -0.0447<br>(0.0785) | 0.0025<br>(0.0689) | 0.0498<br>(0.0982) | -0.0196<br>(0.1055) |
| H5 | -0.0000<br>(0.0095) | -0.0000<br>(0.0109) | -0.0867<br>(0.0956) | -0.3449<br>(0.1618) | -0.0173<br>(0.0158) | -0.0293<br>(0.0184) | -0.0532<br>(0.0482) | -0.2327<br>(0.0295) |

#### 3.2.2 Simulation setting II: variable selection assessment

We examined the performance of using iterative SIS and FDR to select the true mediators **M** from **M**_**0**_. Fig.2 shows the bias of 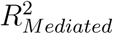 using iterative SIS and FDR to perform variable selection when **M**^(2)^ or **M**^(1)^ were included. The numerical values of the bias, SD, and MSE of the 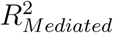 and the product measure estimated by Lasso regression are presented in Table S5. We found that: (1) when only **M**^(1)^ existed, using an inappropriate variable selection method, i.e., FDR, introduced large bias (Fig.2 D); (2) when **M**^(2)^ existed, applying iterative SIS reduced bias to a much smaller scale, while including all variables without variable selection had a large amount of upward bias (Fig.2 A). The FDR method was so conservative in picking up the true mediators, i.e., low true positive rates, that the bias was changed to negative values (Fig.2 B and D). Although not shown, we varied the FDR cutoffs from 0.01 to 0.25 and found that a more liberal cutoff sometimes better controlled the amount of bias, depending on the percentage of true mediators. Nonetheless, the true proportion of mediators is usually unknown. Considering that the FDR method ignores the correlation among the mediators and identifies the true mediators through marginal models, we decided to use iterative SIS for variable selection in the following analyses. We also simulated (V1) by including 13,500 additional noise variables to make this scenario as close as possible to our real-data application for *p*_1_ = 0 and *p*_1_ = 150, and we did not find much difference in terms of the bias, SD, and MSE from our estimation procedure (see Supplementary Materials Section 1.6.1 for details).

**Figure 2.**
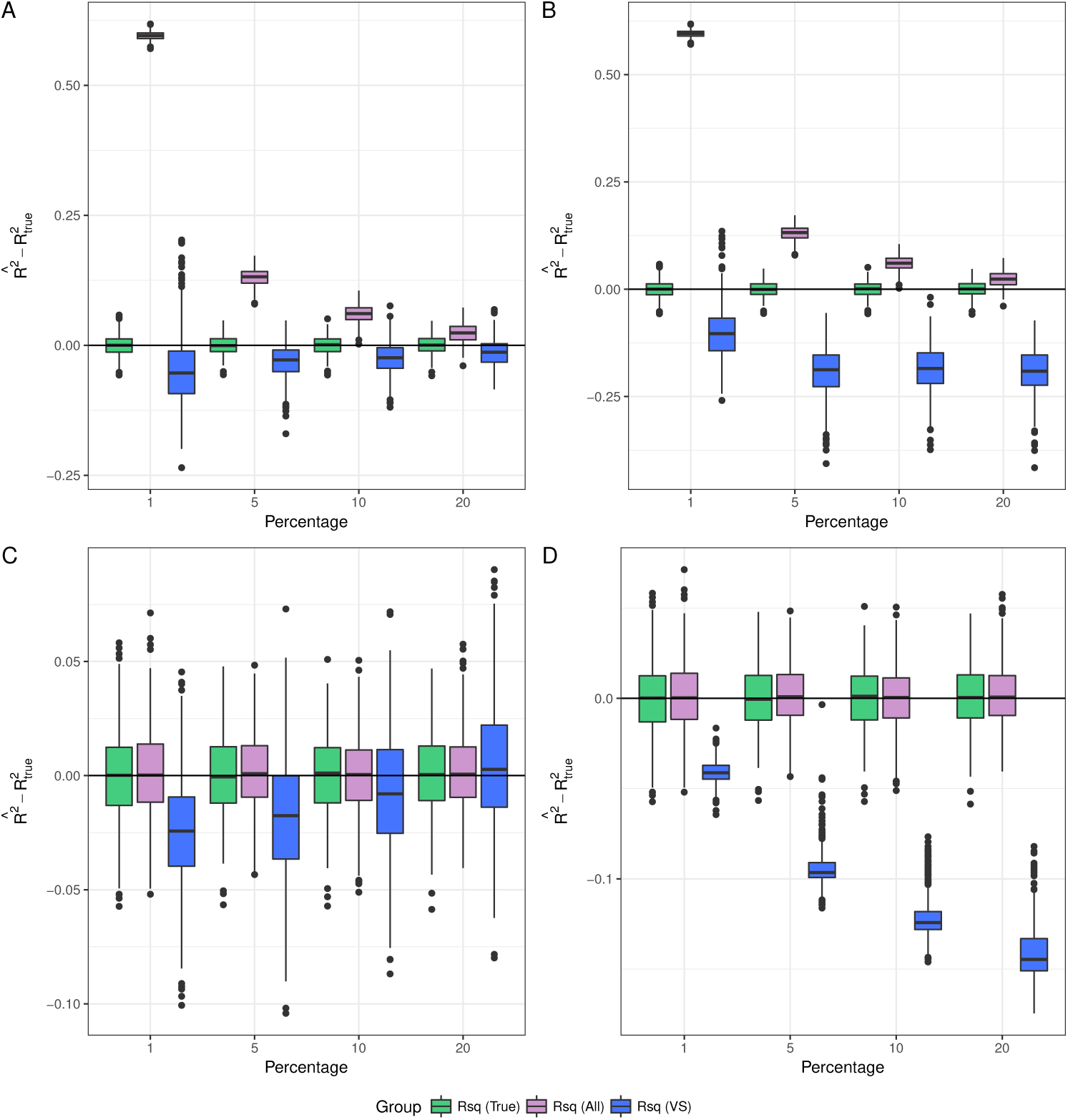
Boxplots of the bias across simulation replications based on a two-step variable selection method, either the iterative SIS or FDR (simulation setting II). X-axis corresponds to the percentage of true mediators; Y-axis corresponds bias across simulation replications. A and B: non-mediators **M**^(2)^ are included in addition to the true mediators; C and D: non-mediators **M**^(1)^ are included in addition to the true mediators. Rsq (All): 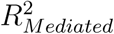 based on all the data without variable selection; Rsq(VS): 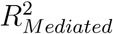 based on the variables selected either by iterative SIS (A and C), or by FDR (B and D); Rsq (True): 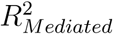 based on the true model/mediator set based on all the data. The numerical values and the bias and variance corresponding to the none mediators (null model) are available in Table S5 and S6.

#### 3.2.3 Finite-sample performance of consistency and CI

We further evaluated the finite-sample performance of the iterative SIS variable selection coupled with the mixed-effect estimation procedure for _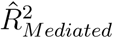. As sample size increased, the bias and SD of 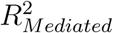 decreased, with a more precise selection of the true mediators (Table S4). In addition, we evaluated the coverage probability of the bootstrap-based CI at different numbers of true mediators (0, 15, 150, and 300) with a sample size of 1500 and found that 100%, 98.0%, 98.0%, and 94.5% CI covered the true value, respectively.

## 4 Real Data Example: the Framingham Heart Study

We hypothesized that the effect of aging on lung function or systolic BP was mediated by changes in GE levels. We performed a mediation analysis on the FHS Offspring Cohort of European ancestry who attended the eighth and ninth examinations with the average interval between visits being around 6 years. Lung function was measured by forced vital capacity (FVC) in liters, using the highest value among at least two acceptable maneuvers. BP was measured as an average of two sequential readings in mmHg. Following Richard et al. (2017), 15 mmHg was added to the systolic BP if a participant reported taking antihypertensive medication at the time of BP measurement. The covariates were the demographic variables of weight in lb, sex, height in inches and smoking status (ever vs never). We focused on subjects with non-missing measurements on the covariates variables, phenotype of interest, and pedigree information, resulting in a final sample size of 1378 for FVC and 1711 for systolic BP. We removed the inter-individual correlation in phenotypes, due to family relatedness, by taking residuals of a linear mixed model with a random effect following a multivariate normal distribution with a zero mean vector and a covariance matrix proportional to the kinship matrix derived from the pedigree information (Cao et al., 2015). GE profiling for 17,873 genes was measured from fasting peripheral whole blood using the Affymetrix Human Exon 1.0 ST GeneChip platform, details of which were described in previous publications (Joehanes et al., 2011). We used age and GE levels at the eighth examination, and FVC and systolic BP at the ninth examination, such that the temporal precedence from exposure to mediators and mediators to phenotype are established.

We assumed that a small proportion of genes were involved in the pathway from aging to the two health traits. As supported by our simulation study (Fig. 2), we did not conduct any pre-screening on **M**^(1)^; instead, we only performed variable selection to exclude **M**^(2)^. The results are summarized in Table 2. We found that the variance in FVC shared by aging and GE was not significantly different from 0, whereas the variance in systolic BP was. Specifically, 19% of FVC variation can be explained by aging, but the number of selected mediators using iterative SIS-MCP was 0 for FVC, suggesting that changes in GE levels did not impact FVC after adjusting for aging. This was further confirmed using the Lasso regression method. A number of GE changes appeared to be associated with aging according to Fig.S3, but we did not observe such a relationship between GEs and FVC. Since GE levels were collected from whole blood, rather than lung tissue, the GE levels in blood might be less relevant for lung function than for blood traits. On the other hand, we found that 6.4% of systolic BP variation can be explained by aging, and 3.3% (95% CI=(0.6%,7.3%)) could be commonly explained by aging and 182 genes whose GEs selected by iterative SIS, accounting for 52% (CI=(12.7%,95.4%)) of the variance explained by aging, as measured by SOS. Note that based on the proportion mediated, 67% (CI=(−119%,175%)) of the total effect was mediated by GEs, whose CI covered 0 and was less interpretable. We further conducted a pathway enrichment analysis of the selected mediators for systolic BP and four nominally significant pathways had biological evidence supporting their potential mediation role between aging and systolic BP (Table S7). For example, the nucleotide excision repair pathway was shown to be involved in age-related vascular dysfunction, which in turn is associated with hypertension (Durik et al., 2012). Finally, according to Fig.S3, the direction of effects through individual pathways were bidirectional, which may explain why the CI of the product, ratio, and proportion measures for systolic BP covered 0. Future analyses with larger sample sizes and using more relevant tissues for lung function are warranted to estimate the total mediation effects.

**Table 2.** Mediation effect size estimated using the Framingham Heart Study data. 95% CI is within the parentheses based on percentiles of 500 bootstrap samples; p is the number of genes whose GE used in estimation; n is the sample size for each trait; A mixed model is used to estimate the quantities, including R^2^’s, ab (the product measure), prop (the proportion mediated), and ratio (ratio mediated).

| Outcome | *n* | *p* | $R^2_{Y,X}$ | $R^2_{Y,M}$ | $R^2_{Mediated}$ | SOS | ab | prop | ratio |
| --- | --- | --- | --- | --- | --- | --- | --- | --- | --- |
| iterative SIS + mixed-effect model |  |  |  |  |  |  |  |  |  |
| FVC <sup>1</sup> | 1378 | 0<br>(0,0) | 0.190<br>(0.144,0.238) | 0<br>(0,0) | 0<br>(0,0) | 0<br>(0,0) | 0<br>(0,0) | 0<br>(0,0) | 0<br>(0,0) |
| Systolic BP | 1711 | 182<br>(147,219) | 0.064<br>(0.033,0.101) | 0.104<br>(0.144,0.324) | 0.033<br>(0.006,0073) | 0.520<br>(0.127,0.954) | 2.28<br>(-3.85,5.91) | 0.67<br>(-1.19,1.76) | 2.08<br>(-15.91,15.85) |

<sup>1</sup> Lasso method was also applied on FVC, by which none of the GE was selected.

## 5 Discussion

We have extended the existing Rsq measure, originally proposed in the single-mediator model, to multiple- and high-dimensional mediator models. Different from the estimation method of the single-mediator model, we proposed a top-down approach: instead of estimating every single regression coefficient, we estimated 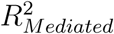 based on the variance components of random coefficients in the mixed model framework. This method can be very efficient, particularly for huge numbers of mediators, because it greatly reduces the number of parameters needed to be estimated. Although 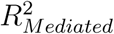 has a good finite-sample performance for models with only true mediators and is robust to the inclusion of certain types of non-mediators, it is biased when variables associated with the exposure, yet not with the dependent variable, are included. To this end, we showed that using iterative SIS can largely mitigate such bias, while using all available GEs led to overestimation, and using a hypothesis testing method with stringent FDR cutoff led to underestimation. To draw valid post-selection inference following the variable selection step, we split the data into halves: we use the first half for variable selection and the second half for estimation. But it is also possible and probably more efficient, though not yet thoroughly studied for iterative SIS, to use all the data (with certain adjustments) in a more unified framework (Lee et al., 2016). We used the nonparametric bootstrap method to calculate the CI and showed that it has satisfactory coverage probability with the sample size comparable to the FHS data.

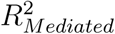 was previously considered to have only a heuristic value, mainly because it can be negative under certain circumstances. When that happens, researchers find it difficult to interpret (Preacher and Kelley, 2011). We emphasize that the 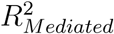 measure is a second-order common effect and thus no longer a proportion measure (Seibold and McPhee, 1979). To facilitate the use of 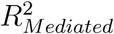, we evaluated the range of the 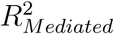 in Supplementary Materials Section 1.2.3 Propositions 1-3. Generally, when the magnitude of the ratio of direct effect and total effect exceeds a certain threshold (larger than 1), 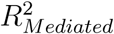 becomes negative; however, under high-dimensional settings, the threshold can be very high. Finally, we have developed an R package ‘RsqMed’, which is publicly available at https://github.com/ytzhong/highD-mediation-analysis, to implement the proposed 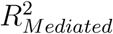 measure estimation and its CI.

## Acknowledgments

This research was supported by the National Institutes of Health (NIH) grants R01CA169122 and R21HL126032; P.W. was supported by NIH grant R01HL116720; H.C. was supported by NIH grants R00HL130593 and U01HL120393. The Framingham Heart Study is conducted and supported by the National Heart, Lung, and Blood Institute (NHLBI) in collaboration with Boston University (Contract No. N01-HC-25195). The authors acknowledge the Texas Advanced Computing Center at The University of Texas at Austin for providing HPC resources. The authors thank Dr. David MacKinnon for discussions in the early stage of this work, and Dr. Lee Ann Chastain and Ms. Jessica Swann for editorial assistance. The authors declare no conflict of interest.

