## Supplementary Materials for "Estimation of Mediation Effect for High-dimensional Omics Mediators with Application to the Framingham Heart Study"

### 1.1 Rsq measure under a single-mediator model

To facilitate the understanding of the Rsq measure, we illustrate it in Fig.S1 via a Venn diagram, in which the area where the three circles overlap is $R^2_{mediated}$. The $R^2_{mediated}$ is interpreted as the variance of Y commonly explained by $X$ and $M$, as in the commonality analysis (CA).

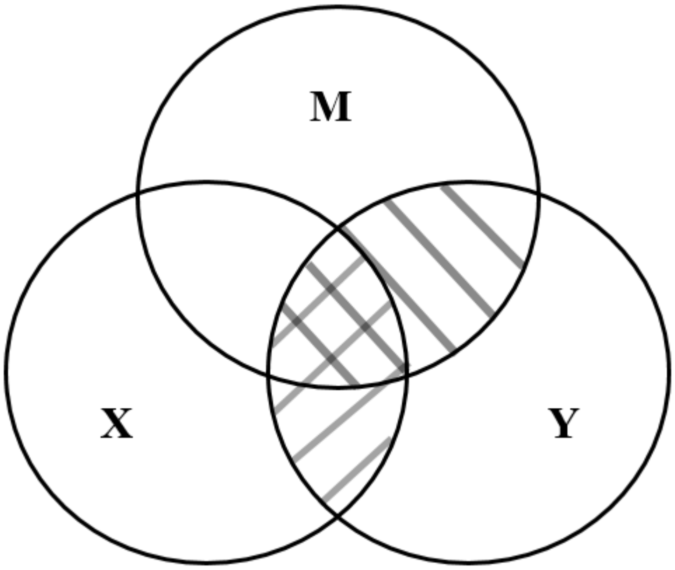


Figure S1: Venn diagram demonstrating the Rsq measure in the single-mediator model: Area indicated with backslashes represents $R^2_{Y,M}$; area with forward slashes represents $R^2_{Y,X}$; and all the covered area represents $R^2_{Y,MX}$. The area in which the three circles overlap represents $R^2_{Mediated}$.

### 1.2 Characteristics of the Rsq measure

The Rsq measure has many characteristics of a good effect size measure, yet it is considered to have heuristic value in certain situations, mainly because of one limitation: it can be negative in the single-mediator model, limiting its interpretation (Preacher and Kelley, 2011). The derivation of the Rsq measure originally came from CA (Fairchild et al., 2009), which is widely used in psychology and education research. CA partitions the regression $R^2$ into unique and common effects. The unique effects indicate that the variance is uniquely accounted for by the independent variable, while the common effects indicate that the variance is common to both the independent variable and the mediator(s). The negativity of the unique and common effects is well accepted in CA, where suppression is considered to be the main reason (Ray-Mukherjee et al., 2014). In fact, it is believed that the suppression brings in extra information that clarifies or "purifies" the relationship between the dependent variable and the independent variable and leads to a practical increment in predictive validity (Ludlow and Klein, 2014).

In high-dimensional settings, suppression is very likely to occur. In light of the interpretation concern, we prove that the Rsq measure is always positive under a consistent model (Proposition 1). We further show that the Rsq measure is always positive under the general consistent model (Proposition 2), where suppression occurs. We find that the negativity is caused by suppression, but suppression does not always lead to the negativity of the Rsq measure. In fact, under the high-dimensional setting, the scenario in which Rsq is negative is unlikely to occur and thus is negligible (Proposition 3). In addition, we did not encounter this concern in our real data analysis. In summary, the Rsq measure is always positive under consistent and general consistent models and is often a trivial case under the high-dimensional setting in which the Rsq measure becomes negative.

Additionally, we investigate the potential bias introduced by two types of non-mediators in Proposition 4 and 5. We also briefly show that our proposed $R^2_{mediated}$ is invariant to certain transformations that may be useful for estimation under the high-dimensional setting in Proposition 6.

#### 1.2.1 $R^2_{Mediated}$ estimand

For exposition purpose, we derive the $R^2_{Mediated}$ under the fixed-effect model with no covariates. Suppose we have the following set of regression models:

$$Y = cX + e_1,$$

$$Y = rX + \sum_{j=1}^{p} M_j b_j + e_2,$$

$$M_j = a_j X + \xi_j,$$

where $p$ is the number of mediators, $e_2 \sim N(0, \phi_1)$, and $\xi = [\xi_1, \xi_2, \ldots, \xi_p] \sim N(0, \mathbf{D}_{p\times p})$. Without loss of generality, X and M are assumed to have variance of 1, and Y is centered at 0. $R^2_{Mediated}$ can be shown to be a function of $\mathbf{a}$ ($[a_1, a_2, \ldots, a_p]^T$), $\mathbf{b}$ ($[b_1, b_2, \ldots, b_p]^T$) and $r$. $R^2_{Mediated}$ is defined as

$$R^2_{Mediated} = R^2_{Y,M} + r^2_{Y,X} - R^2_{Y,MX},$$

where $r_{Y,X} = cor(Y, X)$, $R^2_{Y,MX} = var(rX + \sum_{j=1}^{p} M_j b_j)/\sigma^2_Y$, $\sigma^2_Y$ is the variance of Y, and

$$R^2_{Y,M} = h^T V^{-1}_{MM} h,$$

We have $h = (r_{M_1 Y}, \ldots, r_{M_p Y})^T$ and $V_{MM}$ as a $p \times p$ matrix with $cor(M_i, M_j)$ as the $(i, j)^{th}$ component. By some algebra, it can be shown that

$$R^2_{Y,MX} = \frac{(r + \mathbf{b}^T\mathbf{a})^2 + \mathbf{b}^T\mathbf{D}\mathbf{b}}{\sigma^2_Y}, \tag{S1}$$

$$r^2_{Y,X} = \frac{(r + \mathbf{b}^T\mathbf{a})^2}{\sigma^2_Y}, \tag{S2}$$

$$R^2_{Y,M} = \frac{(r + \mathbf{b}^T\mathbf{a})^2 - r^2/(1 + \mathbf{a}^T\mathbf{D}^{-1}\mathbf{a}) + \mathbf{b}^T\mathbf{D}\mathbf{b}}{\sigma^2_Y}, \tag{S3}$$

such that

$$R^2_{Mediated} = \frac{(r + \mathbf{b}^T\mathbf{a})^2 - r^2/(1 + \mathbf{a}^T\mathbf{D}^{-1}\mathbf{a})}{\sigma^2_Y}, \tag{S4}$$

where $\sigma_Y^2 = (r + \mathbf{b}^T\mathbf{a})^2 + \mathbf{b}^T\mathbf{D}\mathbf{b} + \phi_1$.

#### 1.2.2 Relationships among the mediation effect measures

The relationship between $R^2_{Mediated}$ and the product measure cannot be clearly seen under the mixed-effect model. In the fixed-effect model, $R^2_{Mediated}$ can be expressed in terms of $\mathbf{a}$, $\mathbf{b}$ and $r$ as in equation (S4).

From equation (S4), we can see that the variance of Y explained by M is a function of $a$,$b$ and $r$. Under the circumstances where $r$ is small, the product measure ($\mathbf{b}^T\mathbf{a}$) and $R^2_{Mediated}$ can have high correlation; however, $R^2_{Mediated}$ is not guaranteed to monotonically increase with the product measure. The proportion measure is also a function of $a$,$b$, and $r$, but it decreases as $r$ increases, which is not true for $R^2_{Mediated}$. In fact, the proportion measure has a closer relationship with shared over simple effect (SOS), and SOS can be expressed as

$$\mathrm{SOS} = \frac{R^2_{Mediated}}{R^2_{Y,X}} = 1 - \frac{r^2}{(r + \mathbf{b}^T\mathbf{a})^2(1 + \mathbf{a}\mathbf{D}^{-1}\mathbf{a})}.$$

After some transformation, we can obtain the following equation:

$$1 - \mathrm{proportion}^2 = (1 - \mathrm{SOS})(1 + \mathbf{a}^T\mathbf{D}^{-1}\mathbf{a}).$$

As the proportion measure is positive, the SOS and the proportion measure monotonically increase with each other. When SOS equals 1, the proportion mediated also equals 1. When no mediator exists, both SOS and the proportion mediated equal 0; however, when $\mathbf{b}^T\mathbf{a} = 0$ but some individual pathways do not ($a_jb_j \neq 0$ for some $j$), the proportion mediated cannot capture the mediation effect (proportion$= 0$) while SOS can (SOS $\neq 0$). On the other hand, it is quite rare to happen when SOS equals 0, but proportion mediated does not (r has to be a certain function of a and b, $r = \frac{\mathbf{b}^T\mathbf{a}(1 \pm 1/\sqrt{1+\mathbf{a}^T\mathbf{D}^{-1}\mathbf{a}})}{1 - 1/\mathbf{a}^T\mathbf{D}^{-1}\mathbf{a}}$. We note that $R^2_{Mediated}$ and SOS are both zero when $a_1 = a_2 = \ldots = a_p = 0$; when almost all the variance of Y explained by the exposure is through the mediators, $R^2_{Mediated}$ is close to $R^2_{Y,X}$ and thus SOS $\approx 1$; $R^2_{Mediated}$ and SOS approach 1 when $\mathbf{b}^T\mathbf{a}$ goes to infinity.

#### 1.2.3 Range of the Rsq measure

**Proposition 1**: In the consistent model, where $a_j b_j$ and $r$ are in the same direction ($a_j b_j r > 0$) for $j = 1, 2, \ldots, p$, $R^2_{Mediated} \in (0, 1)$.

Proof: To show $R^2_{Mediated} > 0$ is equivalent to showing that the nominator $(r + \mathbf{b}^T\mathbf{a})^2 - r^2/(1 + \mathbf{a}^T\mathbf{D}^{-1}\mathbf{a}) > 0$.

Since $D$ is a (semi)positive definite matrix, $\mathbf{a}^T\mathbf{D}^{-1}\mathbf{a} > 0$.

Thus, $(r + \mathbf{b}^T\mathbf{a})^2 - r^2/(1 + \mathbf{a}^T\mathbf{D}^{-1}\mathbf{a}) > (\mathbf{b}^T\mathbf{a})^2 + 2r\mathbf{b}^T\mathbf{a} > 0$.

In addition, $(r + \mathbf{b}^T\mathbf{a})^2 - r^2/(1 + \mathbf{a}^T\mathbf{D}^{-1}\mathbf{a}) < (r + \mathbf{b}^T\mathbf{a})^2 + \mathbf{b}^T\mathbf{D}\mathbf{b} + \tau$ for any $\mathbf{a}$, $\mathbf{b}$ and $r$.

When $\mathbf{b}^T\mathbf{a} \to \infty$, $R^2_{Mediated} \to 1$.

Therefore, $R^2_{Mediated} \in (0, 1)$ under the consistent model.

Next, we extend the proof to scenarios where $\mathbf{b}^T\mathbf{a}$ and $r$ are in the same direction ($\mathbf{b}^T\mathbf{a}r > 0$), and we define such models herein as general consistent models.

**Proposition 2**: In the general consistent model, $R^2_{Mediated} > 0$.

Proof: Under the general consistent model, $(r + \mathbf{b}^T\mathbf{a})^2 > r^2$.

In addition, since $D$ is a positive (semi)definite matrix, $r^2 > r^2/(1 + \mathbf{a}^T\mathbf{D}^{-1}\mathbf{a})$.

Therefore, $(r + \mathbf{b}^T\mathbf{a})^2 > r^2 > r^2/(1 + \mathbf{a}^T\mathbf{D}^{-1}\mathbf{a})$ . Since $\sigma^2_Y > 0$, $R^2_{Mediated}$ is always positive.

**Proposition 3**: When $|r/c| > \sqrt{1 + \mathbf{a}^T\mathbf{D}^{-1}\mathbf{a}}$, $R^2_{Mediated}$ is negative.

Proof: When $r$ is large and $c \approx 0$,

$$R^2_{Mediated} \approx -\frac{r^2/(1 + \mathbf{a}^T\mathbf{D}^{-1}\mathbf{a})}{\sigma^2_Y}.$$

Since $r^2 > 0$ and $1 + \mathbf{a}^T\mathbf{D}^{-1}\mathbf{a} > 0$, $R^2_{Mediated} < 0$.

The first step to establish mediation according to Baron and Kenny (1986) is that the independent variable must affect the dependent variable, that is, the total effect $c$ should be different from 0. If the effect is not significant, the analysis for consistent mediation stops. Coinciding with this step, we show that the negativity of the Rsq measure happens when this criterion does not hold.

More generally, it can be proven that $R^2_{Mediated} < 0$ when $|r/c| > \sqrt{1 + \mathbf{a}^T\mathbf{D}^{-1}\mathbf{a}}$ using algebra. Under the high-dimensional setting, $\mathbf{a}^T\mathbf{D}^{-1}\mathbf{a}$ can be large and $c$ is large enough to pass the first step of Baron and Kenny (1986); therefore, the scenario where $R^2_{Mediated} < 0$ may not be very likely to happen.

#### 1.2.4 Non-mediators and transformation

Variables that are not mediators may be falsely included in the mediation model. Here, we denote three types of non-mediators: $\mathbf{M}^{(1)} = \{M_j : b_j \neq 0, a_j = 0\}$ as the set of variables $M^{(1)}$, $\mathbf{M}^{(2)} = \{M_j : b_j = 0, a_j \neq 0\}$ as the set of variables $M^{(2)}$, and noise variables as the set $\{M_j : b_j = 0, a_j = 0\}$. Let $\hat{\mathbf{M}}$ be the set of variables included in the model, $\mathbf{M} = \{M_j : b_j \neq 0, a_j \neq 0\}$ be the true mediator set consisting of a total of $t$ true mediating variables.

**Proposition 4**: $\mathbf{M^{(1)}}$ in the mediation model does not bias the estimation of $R^2_{Mediated}$.

Proof: Without loss of generality, we assume $\hat{\mathbf{M}} = [\mathbf{M}, \mathbf{M^{(1)}}]$, then $\mathbf{a}^T = [a_1, a_2, \ldots, a_t, 0, \ldots, 0]$. By some linear algebra, it can be shown that $\mathbf{a}^T\mathbf{D}^{-1}\mathbf{a} = [a_1, a_2, \ldots, a_t]\mathbf{D_{tt}}[a_1, a_2, \ldots, a_t]^T$, where $\mathbf{D_{tt}}$ is $\mathbf{D}^{-1}$ with the first t columns and the first t rows.

In addition, $\mathbf{b}^T\mathbf{a} = [b_1, b_2, \ldots, b_t][a_1, a_2, \ldots, a_t]^T$. Therefore,

$$R^2_{Mediated}(\hat{\mathbf{M}}) = R^2_{Mediated}(\mathbf{M})$$

In the special case where all $\mathbf{M} = \emptyset$,

$$R^2_{Y,MX} = R^2_{Y,M} + r^2_{Y,X},$$

$$R^2_{Mediated} = R^2_{Y,M} + r^2_{Y,X} - (R^2_{Y,M} + r^2_{Y,X}) = 0.$$

Therefore, the inclusion of $\mathbf{M^{(1)}}$ does not lead to bias in the point estimate of $R^2_{Mediated}$. When such variables are included in the mediation model, both $R^2_{Y,M}$ and $R^2_{Y,XM}$ increase the same amount. As a result, their effects cancel out in $R^2_{Mediated}$. Since the noise variables also have $a_j = 0$, they do not bias the estimation either.

**Proposition 5**: $\mathbf{M^{(2)}}$ in the mediation model bias the estimation of $R^2_{Mediated}$.

Proof: Without loss of generality, we let $\hat{\mathbf{M}} = [\mathbf{M}, \mathbf{M}^{(\mathbf{2})}]$, then $\mathbf{a}^T = [a_1, a_2, \ldots, a_t, a'_1, \ldots, a'_q]$, $\mathbf{b}^T = [b_1, b_2, \ldots, b_t, 0, \ldots, 0]$. In addition, we let

$$\mathbf{D} = \begin{bmatrix} \mathbf{D}_{tt} & \mathbf{D}_{tq} \\ \mathbf{D}_{qt} & \mathbf{D}_{qq} \end{bmatrix}$$

By some linear algebra, it can be shown that

$$\begin{aligned} \mathbf{a}^T\mathbf{D}^{-1}\mathbf{a} = [a_1, a_2, \ldots, a_t]\mathbf{D_{tt}}[a_1, a_2, \ldots, a_t]^T + 2[a_1, a_2, \ldots, a_t]\mathbf{D_{tq}}[a'_1, a'_2, \ldots, a'_q]^T \\ + [a'_1, a'_2, \ldots, a'_q]\mathbf{D_{qq}}[a'_1, a'_2, \ldots, a'_q]^T \\ \neq [a_1, a_2, \ldots, a_t]\mathbf{D_{tt}}[a_1, a_2, \ldots, a_t]^T. \end{aligned}$$

Thus $R^2_{Y,M}$ is biased away from its true value. In addition, since $\mathbf{b}^T\mathbf{a} = [b_1, b_2, \ldots, b_t][a_1, a_2, \ldots, a_t]^T$, $R^2_{Y,MX}$ and $R^2_{Y,X}$ stay the same. We conclude that

$$R^2_{Mediated}(\hat{\mathbf{M}}) \neq R^2_{Mediated}(\mathbf{M}).$$

Therefore, inclusion of $\mathbf{M}^{(\mathbf{2})}$ is problematic.

Transforming correlated mediators in the original model to be uncorrelated may be helpful in dimension reduction, especially for high-dimensional data (Huang and Pan, 2016).

**Proposition 6**: $R^2_{Mediated}$ is invariant to the following transformation of mediators: the mediators are transformed from $\mathbf{M}$ to $\mathbf{P}$: $\mathbf{P} = \mathbf{uM}$, $\mathbf{u}$ is an orthogonal invertible $p \times p$ matrix (Huang and Pan, 2016).

Proof: $\mathbf{P}_i = i_3^* + X_i\mathbf{a}^* + \xi_i^*$, where $\mathbf{a}^* = \mathbf{ua}$ and $\xi_i^* = \mathbf{u}\xi_i$. $\xi_i \sim N(0, \mathbf{D}^*)$ with $\mathbf{D}^* = \mathbf{uDu}^T$. Therefore, $\mathbf{a}^{*T}\mathbf{D}^*\mathbf{a}^* = \mathbf{a}^T\mathbf{Da}$.

Similarly, $Y = i_2 + rX + \mathbf{M}^T\mathbf{b} + e_2 = i_2 + rX + \mathbf{P}^T\mathbf{b}^* + e_2$, where $\mathbf{b}^* = (\mathbf{u}^T)^{-1}\mathbf{b}$. Therefore, $\mathbf{b}^{*T}\mathbf{a}^* = \mathbf{b}^T\mathbf{a}$.

In conclusion, such a transformation does not influence $R^2_{Mediated}$. Possible transformations include $\mathbf{u}$ such that $\mathbf{uDu}^T = diag(\mathbf{D})$.

#### 1.2.5 Consistency

**Proposition 6**: The estimation procedure coupled with iterative SIS-based variable selection guarantees the consistency of $R^2_{Mediated}$.

Proof: Let the putative mediation variables in the initial assessment as $\mathbf{M}_0$, $\mathbf{A} = \{j : b_j \neq 0\}$ be the set of indices of $\mathbf{M}_0$. $\mathbf{A}_1^* = \{j : \hat{b}_j^M \neq 0\}$, where $\hat{b}_j^M$ is the estimated marginal correlation coefficient between the $j^{th}$ mediator and the dependent variable. $\mathbf{A}_2^* = \{j : \hat{b}_j^{MCP} \neq 0\}$, where $\hat{b}_j^{MCP}$ is the estimated regression coefficient under the minimax concave penalty (MCP).

It has been proven that correlation learning in the sure independence screening (SIS) method has the sure screening property, that is, $P(\mathbf{A} \subset \mathbf{A}_1^*) \to 1$ as $n \to \infty$ under mild conditions (Fan and Lv, 2008). MCP is applied after the first dimension reduction step. Since it has the oracle property, for $\forall j \in \mathbf{A}$, $P(j \in \mathbf{A}_2^*) \to 1$ and $\forall j' \notin \mathbf{A}$, $P(j' \in \mathbf{A}_2^*) \to 0$ as $n \to \infty$.

The $R^2_{Mediated} = R^2_{Y,M} + r^2_{Y,X} - R^2_{Y,MX}$, and each component is estimated in the second half of the data as follows:

$$\hat{R}^2_{Y,MX} = 1 - \frac{\hat{\phi}_1}{\hat{\sigma}^2_Y}, \hat{R}^2_{Y,M} = 1 - \frac{\hat{\phi}_2}{\hat{\sigma}^2_Y}, \hat{r}^2_{Y,X} = \frac{\hat{c}^2}{\hat{\sigma}^2_Y},$$

where $\hat{c}$ is the MLE of the total effect; $\hat{\sigma}^2_Y = \sum_{i=1}^n (Y^2 - \bar{Y})/(n-1)$; $\hat{c}$ and $\hat{\sigma}^2_Y$ are consistent estimates of $c$ and $\sigma^2$. Moreover, we estimate $\hat{\phi}_1$ and $\hat{\phi}_2$ by a mixed-effect model using REML, which is consistent under mild conditions (Cressie and Lahiri, 1993). According to the continuous mapping theorem, $\hat{R}^2_{Mediated}$ is a consistent estimate of $R^2_{Mediated}$: $\hat{R}^2_{Mediated} \xrightarrow{p} R^2_{Mediated} | \hat{\mathbf{A}} = \mathbf{A}$. That is, for every $\varepsilon > 0$, $\lim_{n\to\infty} P(|\hat{R}^2_{Mediated}(n) - R^2_{Mediated}| > \varepsilon | \hat{\mathbf{A}} = \mathbf{A}) = 0$.

Therefore,

$$\begin{aligned}
\lim_{n\to\infty} P(|\hat{R}^2_{Mediated}(n) - R^2_{Mediated}| > \varepsilon|) &= \lim_{n\to\infty} P(|\hat{R}^2_{Mediated}(n) - R^2_{Mediated}| > \varepsilon | \hat{\mathbf{A}} = \mathbf{A}) P(\hat{\mathbf{A}} = \mathbf{A}) \\
&+ \lim_{n\to\infty} P(|\hat{R}^2_{Mediated}(n) - R^2_{Mediated}| > \varepsilon | \hat{\mathbf{A}} \neq \mathbf{A}) P(\hat{\mathbf{A}} \neq \mathbf{A}) \\
&= 0 \times 1 + \lim_{n\to\infty} P(|\hat{R}^2_{Mediated}(n) - R^2_{Mediated}| > \varepsilon | \hat{\mathbf{A}} \neq \mathbf{A}) \times 0 \\
&= 0.
\end{aligned}$$

In conclusion, the consistency of $R^2_{Mediated}$ is established for using SIS-MCP to perform variable selection and the mixed-effect model to estimate the variance components. The proof holds for any SIS with oracle property; iterative SIS served as an extension to the SIS-based model selection method where the regularity conditions may fail. It has been proven that iterative SIS also has the sure screening property (Fan et al., 2009). Therefore, the result holds for our proposed method based on iterative SIS-MCP as well.

### 1.3 Rsq measure under a model with a directed correlation structure among mediators

When there is a directed correlation structure among the mediators, the proposed Rsq measure is still a valid effect size measure to capture the variation of the dependent variable explained by the independent variable through the mediators. For example, if there are two mediators $M_1$ and $M_2$ from independent $X$ to outcome variable $Y$, $M_1$ can affect $M_2$. There are four pathways from $X$ to $Y$, that is $X - M_1 - M_2 - Y$, $X - M_1 - Y$, $X - M_2 - Y$ and $X - Y$. We can write the corresponding models as:

$$Y = i_1 + cX + e_1,$$

$$Y = i_2 + rX + M_1 b_1 + M_2 b_2 + e_2,$$

$$M_1 = i_{31} + a_1 X + \xi_1, \tag{S5}$$

$$M_2 = i_{32} + a_2' X + \gamma M_1 + \xi_2'. \tag{S6}$$

Model (S6) can be further written as the following form by plugging in equation (S5):

$$M_2 = i_{32} + (a_2' + \gamma a_1)X + (\gamma \xi_1 + \xi_2') = i_{32} + a_2 X + \xi_2.$$

We denote $a_2 = (a_2' + \gamma a_1)$ and $\xi_2 = (\gamma \xi_1 + \xi_2')$. VanderWeele and Vansteelandt (2014) showed that the product measure under this framework is $a_1 b_1 + a_2 b_2$. For our Rsq measure, the residuals $\xi_1$ and $\xi_2$ are correlated and the corresponding correlation matrix $\mathbf{D}_{p\times p}$ is no longer diagonal. However, equations (S1), (S2), (S3) and (S4) still hold. The complicated causal structure is reflected and captured by $\mathbf{D}_{p\times p}$ matrix. We have performed a simulation study to assess bias under the low-dimensional setting, as demonstrated in Supplementary Material Section 1.5.1.

### 1.4 Bootstrap-based confidence interval for $R^2_{Mediated}$

When variable selection is involved, the point estimate of $R^2_{Mediated}$ can be calculated through the following steps in general:

1. Randomly split the data into two non-overlapping parts. Regress the dependent variable on the covariates to obtain the residuals in the first part and the second part;
2. Using the residuals as the 'adjusted' dependent variable to build an iterative SIS model for mediator selection in the first part;
3. Standardize the selected mediators and obtain the corresponding point estimate of $R^2_{Mediated}$ in the second part following the main text Section 2.3.1.

To obtain the CI of the $R^2_{Mediated}$, we propose using the following bootstrap-based procedure:

A. Draw B bootstrap samples of size $n$ with replacement from the original sample;
B. Calculate the point estimate of $R^2_{Mediated}$ for each bootstrapped sample based on the above iterative SIS-mixed model procedure 1-3, denoted as $\hat{R}^2_1, \hat{R}^2_2, \ldots, \hat{R}^2_B$;
C. Extract the 2.5% and 97.5% quantiles of $\hat{R}^2_1, \hat{R}^2_2, \ldots, \hat{R}^2_B$ as the lower and upper bound of CI.

We have developed an R package 'RsqMed' for researchers and practitioners to estimate the Rsq measure and its CI based on the above described procedure. The package is publicly available at the GitHub website https://github.com/ytzhong/highD-mediation-analysis. We would recommend implementing the iterative SIS model selection procedure with the BIC to select the tunning parameter. Variable selection methods based on cross-validation are not supported in the current version of implementation; however, an extension is feasible but has to be done with caution (non-overlapping validation and training samples are required to avoid overfitting, which is not supported by the standard bootstrap procedure). Other resampling methods to correct post-selection bias are also possible (Sun and Bull, 2005).

### 1.5 Additional Simulations

#### 1.5.1 Bias and variance under the low-dimensional setting

We assessed the bias and variance of different types of mediation effect size measures under the low-dimensional setting. Data were generated using model (2) and model (3). The sample size was $n = 1500$, $e_2 \sim N(0,1), X \sim N(0,1)$ and $r = 1$. There were $p$ variables in $\mathbf{M_0}$, and $\xi = [\xi_1, \xi_2, \ldots, \xi_p] \sim N(0, \mathbf{D}_{p\times p})$. We evaluated the performance under the following six low-dimensional scenarios. (L1) and (L5) represent the scenarios of correct model specification, (L2), (L3), and (L4) represent the scenarios of including the three types of non-mediators. In (L1) to (L5), $\mathbf{D}_{p\times p}$ is the identity matrix, mimicking the undirected relationship among mediators. (L6) represents a scenario of a model with directed causal structure among mediators) and $\mathbf{D}_{p\times p}$ has off-diagonal items.

(L1) Two true mediators with same direction were included: $p = 2, a_1 = 1, a_2 = 0.5, b_1 = b_2 = 0.5$;

(L2) Two $M^{(2)}$ were added to L1): $p = 4, a_3 = a_4 = 2, b_3 = b_4 = 0$;

(L3) Two $M^{(1)}$ were added to L1): $p = 4, a_3 = a_4 = 0, b_3 = b_4 = 0.5$;

(L4) Two noise variables were added to L1): $p = 4, a_3 = a_4 = b_3 = b_4 = 0$;

(L5) Two true mediators with different directions were included: $p = 2, a_1 = 1, a_2 = 0.5, b_1 = 0.5, b_2 = -0.5$;

(L6) Mediators followed a correlated causal structure, as shown in Fig.S2 with parameters listed below.

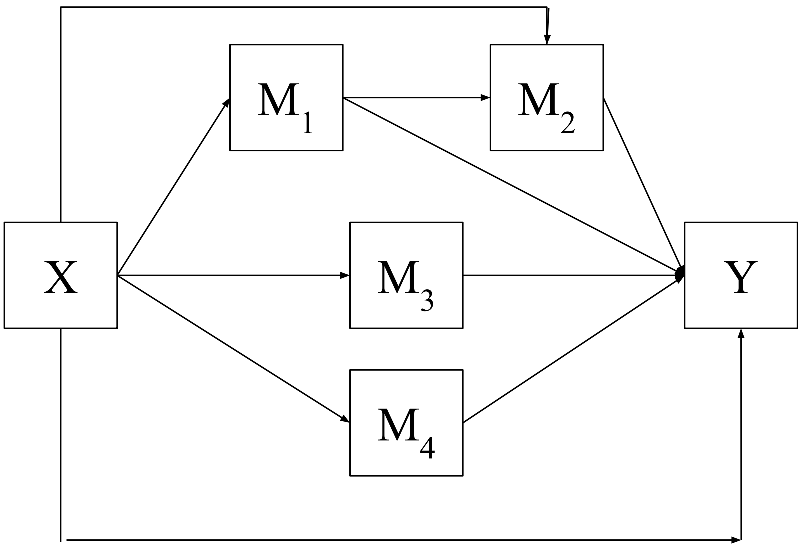


Figure S2: Structural Diagram for simulation setting (L6)

$$E(Y) = X + 0.5M_1 + 0.2M_2 + 0.5M_3 + 0.1M_4,$$
$$E(M_1) = 0.5X, E(M_2) = 0.5X + M_1,$$
$$E(M_3) = 0.5X, E(M_4) = X.$$

Table S1 shows the results of bias and standard deviation assessment under the low-dimensional setting, i.e., (L1) to (L6). The estimators of all the mediation effect size measures had acceptable bias under the six scenarios, except that the $R^2_{Mediated}$ estimators and their closely related SOS estimators were biased under scenario (L2), as expected. In addition, we showed that the bias was small when directed causal structure among the mediators existed in (L6). Overall, $R^2_{Mediated}$ estimated by the mixed-effect model had smaller bias and variance than that estimated based on the fixed-effect model, which also held for SOS. In contrast, using the fixed-effect model gave smaller bias than using the mixed-effect model for the product, ratio, and proportion measures under the low-dimensional setting.

Table S1: Bias and standard deviation under the low-dimensional setting (Simulation setting I): the first row is the bias and the second row is the corresponding standard deviation for each scenario. True values are marked in boldface. (Fixed) indicates that the estimation is based on a fixed-effect model; otherwise, it is estimated by a mixed-effect model.

| | L1 | L2 | L3 | L4 | L5 | L6 |
|---|---|---|---|---|---|---|
| $\mathbf{R^2_{Mediated}}$ | **0.5738** | **0.5738** | **0.5171** | **0.5738** | **0.3651** | **0.7632** |
| $R^2_{Mediated}$ | −0.0002 | 0.0758 | 0.0002 | −0.0003 | −0.0010 | 0.0000 |
| | (0.0141) | (0.0142) | (0.0171) | (0.0146) | (0.0189) | (0.0105) |
| $R^2_{Mediated}$ (Fixed) | −0.0004 | 0.1320 | 0.0006 | −0.0004 | −0.0023 | 0.0004 |
| | (0.0246) | (0.0248) | (0.0330) | (0.0254) | (0.0517) | (0.0177) |
| **SOS** | **0.8549** | **0.8549** | **0.8549** | **0.8549** | **0.7156** | **0.8303** |
| SOS | −0.0010 | 0.1126 | −0.0008 | −0.0008 | −0.0014 | −0.0003 |
| | (0.0125) | (0.0055) | (0.0126) | (0.0120) | (0.0229) | (0.0040) |
| SOS (Fixed) | −0.0011 | 0.1125 | −0.0007 | −0.0006 | −0.0010 | −0.0003 |
| | (0.0125) | (0.0055) | (0.0126) | (0.0120) | (0.0228) | (0.0040) |
| **Product** | **0.75** | **0.75** | **0.75** | **0.75** | **0.25** | **1.95** |
| Product | −0.0052 | −0.0147 | −0.0047 | −0.0064 | −0.0037 | −0.0189 |
| | (0.0349) | (0.0815) | (0.0395) | (0.0337) | (0.0341) | (0.0830) |
| Product (Fixed) | −0.0031 | −0.0106 | −0.0020 | −0.0020 | −0.0015 | −0.0089 |
| | (0.0349) | (0.0826) | (0.0395) | (0.0336) | (0.0343) | (0.0827) |
| **Ratio** | **0.75** | **0.75** | **0.75** | **0.75** | **0.25** | **1.95** |
| Ratio | −0.0061 | −0.0138 | −0.0058 | −0.0087 | −0.0034 | −0.0240 |
| | (0.0584) | (0.1391) | (0.0606) | (0.0567) | (0.0412) | (0.3777) |
| Ratio (Fixed) | −0.0023 | −0.0063 | −0.0011 | −0.0011 | −0.0006 | 0.0226 |
| | (0.0586) | (0.1421) | (0.0610) | (0.0570) | (0.0415) | (0.4055) |
| **Proportion** | **0.4286** | **0.4286** | **0.4286** | **0.4286** | **0.20** | **0.6610** |
| Proportion | −0.0026 | −0.0082 | −0.0026 | −0.0035 | −0.0030 | −0.0068 |
| | (0.0192) | (0.0463) | (0.0200) | (0.0187) | (0.0265) | (0.0294) |
| Proportion (Fixed) | −0.0019 | −0.0069 | −0.0017 | −0.0020 | −0.0016 | −0.0058 |
| | (0.0192) | (0.0466) | (0.0199) | (0.0186) | (0.0265) | (0.0292) |

#### 1.5.2 Bias and variance under the high-dimensional setting

In the main text Table 1, we presented the bias and standard deviation of five different measures, i.e., product, proportion, ratio, $R^2_{Mediated}$, and SOS under the high-dimensional setting. However, they are not directly comparable because they are on different scales. Table S2 presents their true values. Note that the true proportion mediated is negative due to different directions of the mediation effects and the direct effect.

Table S2: True values of the five measures for Table 1 in the main text

| | $R^2_{Mediated}$ | SOS | Product | Proportion | Ratio |
|---|---|---|---|---|---|
| H1 | 0.3968 | 0.8098 | -0.3123 | -0.1162 | -0.1041 |
| H2 | 0.3968 | 0.8098 | -0.3123 | -0.1162 | -0.1041 |
| H3 | 0.0874 | 0.8098 | -0.3123 | -0.1162 | -0.1041 |
| H4 | 0.3968 | 0.8098 | -0.3123 | -0.1162 | -0.1041 |
| H5 | 0.8444 | 0.9723 | 4.0431 | 0.5741 | 1.3477 |

#### 1.5.3 Bias and variance under the high-dimensional setting with model misspecification

We evaluated the robustness of the normal assumption of the random effects under the high-dimensional setting. Normally distributed random effect is commonly seen in the literature. It has been shown to be quite robust in real data analysis and highly correlated data in simulations (Yang et al., 2010, 2019). There are multiple theoretical studies that consistently show the negligible impact of misspecifying random effect distributions for the linear mixed model (Verbeke and Lesaffre, 1997; McCulloch and Neuhaus, 2011; Neuhaus et al., 2013). In the current context, we further evaluated whether misspecifying the distribution of the random effects would bias $R^2_{mediated}$ in the scenarios (H6) to (H10) whose settings are exactly the same as (H1) to (H5) except that $a_j$ and $b_j$ follow a scaled t-distribution with the degree of freedom equal to one. Table S3 confirms that the bias under (H6) to (H10) was not larger than the bias under (H1) to (H5) with correctly-specified random effect distribution.

Table S3: Bias and standard deviation under the high-dimensional setting with model-misspecification (Simulation setting I): the first row is the bias and the second row is the corresponding standard deviation for each scenario. The true values are marked in boldface.

| | H6 | H7 | H8 | H9 | H10 |
|---|---|---|---|---|---|
| $\mathbf{R^2_{Mediated}}$ | **0.2126** | **0.2126** | **0.0280** | **0.2126** | **0.3287** |
| $R^2_{Mediated}$ | 0.0007 | 0.0328 | 0.0005 | 0.0026 | −0.0006 |
| | (0.0206) | (0.0199) | (0.0088) | (0.0207) | (0.0197) |
| **SOS** | **0.8634** | **0.8634** | **0.8634** | **0.8634** | **0.9198** |
| SOS | 0.0028 | 0.1334 | 0.0147 | 0.0106 | −0.0018 |
| | (0.0835) | (0.0807) | (0.2710) | (0.0839) | (0.0551) |

#### 1.5.4 Evaluation of finite-sample performance of consistency

Although we showed the consistency property of our proposed estimation procedure in theory, we are interested in its finite-sample performance. We evaluated the following simulation settings (C1 to C4), and we looked at three sets of sample size $n = 750, 1500$, and 3000 with the initial size of $\mathbf{M_0}$ as $p = 1500$; Additional parameters were the same as V1): $r = 3$, $e_2 \sim N(0, 1)$, and $X \sim N(0, 1)$; $\mathbf{D}_{p\times p}$ is the identity matrix. Four scenarios were assessed: (C1) and (C2) represent different sparsity models when $\mathbf{M}^{(2)}$ are included, (C3) represents when $\mathbf{M}^{(1)}$ and noise variables are included, and (C4) represents when all three types of non-mediators are included:

(C1) There were 15 true mediators and 1485 $\mathbf{M}^{(2)}$: $a_j \sim N(0, 0.2)$ for $j = 1, \ldots, 1500$, and $b_j \sim N(0, 0.2)$ for $j = 1, \ldots, 15$, $b_j = 0$ for $j = 16, \ldots, 1500$.

(C2) There were 150 true mediators, and 1350 $\mathbf{M}^{(2)}$: $a_j \sim N(0, 0.2)$ for $j = 1, \ldots, 1500$, and $b_j \sim N(0, 0.2)$ for $j = 1, \ldots, p_1$, $b_j = 150$ for $j = 151, \ldots, 1500$.

(C3) There were 150 true mediators, 150 $\mathbf{M}^{(1)}$, and 1200 noise variables: $b_j \sim N(0, 0.2)$ for $j = 1, \ldots, 300$ and $b_j = 0$ for $j = 301, \ldots, 1500$; $a_j \sim N(0, 0.2)$ for $j = 1, \ldots, 150$, $a_j = 0$ for $j = 151, \ldots, 1500$.

(C4) There were 150 true mediators, 150 $\mathbf{M}^{(1)}$, 150 $\mathbf{M}^{(2)}$, and 1050 noise variables: $b_j \sim N(0, 0.2)$ for $j = 1, \ldots, 300$, and $b_j = 0$ for $j = 301, \ldots, 1500$; $a_j \sim N(0, 0.2)$ for $j = 1, \ldots, 150, 1200, \ldots, 1500$, $a_j = 0$ for $j = 151, \ldots, 300$.

Table S4: Finite-sample performance of consistency of $R^2_{Mediated}$: the bias, standard deviation (SD), and mean square error (MSE) in $\hat{R}^2_{Mediated}$ decrease with increasing total sample sizes. Initial number of variables under investigation is 1500; $\bar{p}$ is the average number of selected variables; $\bar{p}_M/\bar{p}_{M^{(1)}}/\bar{p}_{M^{(2)}}$ are the average number of selected true mediators, $\mathbf{M}^{(1)}$, and $\mathbf{M}^{(2)}$, respectively; TP: average true positive rate; FP: average false positive rate.

| Sample Size | bias | SD | MSE | $\bar{p}(\bar{p}_M/\bar{p}_{M^{(1)}}/\bar{p}_{M^{(2)}})$ | TP | FP |
|---|---|---|---|---|---|---|
| C1) | | | | | | |
| 750 | -0.0339 | 0.1456 | 0.0224 | 16(7/0/9) | 0.47 | 0.0060 |
| 1500 | -0.0484 | 0.0678 | 0.0069 | 12(10/0/2) | 0.63 | 0.0019 |
| 3000 | -0.0174 | 0.0490 | 0.0027 | 14(12/0/2) | 0.78 | 0.0015 |
| C2) | | | | | | |
| 750 | -0.1029 | 0.0496 | 0.0130 | 61(37/0/24) | 0.24 | 0.0177 |
| 1500 | -0.0253 | 0.0307 | 0.0016 | 128(82/0/46) | 0.54 | 0.0341 |
| 3000 | -0.0229 | 0.0179 | 0.0008 | 123(102/0/21) | 0.68 | 0.0154 |
| C3) | | | | | | |
| 750 | -0.1874 | 0.0637 | 0.0392 | 84 (19/22/0) | 0.14 | 0.036 |
| 1500 | -0.0540 | 0.0339 | 0.0041 | 213 (64/71/0) | 0.45 | 0.064 |
| 3000 | -0.0235 | 0.0205 | 0.0010 | 250 (102/108/0) | 0.71 | 0.033 |
| C4) | | | | | | |
| 750 | -0.1612 | 0.0614 | 0.0298 | 84 (19/24/4) | 0.14 | 0.035 |
| 1500 | -0.0459 | 0.0324 | 0.0032 | 213 (65/71/9) | 0.45 | 0.063 |
| 3000 | -0.0216 | 0.0206 | 0.0009 | 247 (102/106/4) | 0.69 | 0.032 |

#### 1.5.5 Coverage Probability of the bootstrap procedure

To evaluate the coverage probability of the bootstrap-based CI for $R^2_{Mediated}$, we adapted the simulation setting in the first scenario (V1), but let $\mathbf{a}$ and $\mathbf{b}$ vary across 200 replications, i.e., $a_{jd} \sim N(0, 0.2)$ for $j = 1, \ldots, 1500$, where $d$ was the index for replicates and $d = 1, \ldots, 200$; $b_{jd} \sim N(0, 0.2)$ for $j = 1, \ldots, p_1$ and $d = 1, \ldots, 200$, $b_j = 0$ for $j = p_1 + 1, \ldots, 1500$. To mimic the real-data application, we evaluated the coverage probability when the number of true mediators was fixed at $p_1 = 0, 15, 150$ and 300, and sample size at 1500. The coverage probability was 100%, 98.0%, 98.0%, and 94.5%, respectively.

### 1.6 Simulation setting II

The MSE, bias, and standard deviation are presented in Table S5 and S6, corresponding to Figure 2 A and B in the main text. FDR was calculated using the Benjamini and Hochberg method (Benjamini and Hochberg, 1995).

Table S5: The mean squared error (MSE), bias and standard deviation (SD) of $R^2_{Mediated}$ in simulation setting II (V1). Rsq (True) is $R^2_{Mediated}$ based on the true mediators $\mathbf{M}$; Rsq(All) is $R^2_{Mediated}$ based on the putative mediation variables in the initial assessment $\mathbf{M}_0$; Rsq (VS=iSIS-MCP) is $R^2_{Mediated}$ based on selected mediator set $\hat{\mathbf{M}}$ using iterative SIS; Rsq (VS=FDR) is $R^2_{Mediated}$ based on $\hat{\mathbf{M}}$ using the FDR method; ab (Lasso): product measure estimated using Lasso regression for variable selection and estimation simultaneously.

| Percentage | Rsq (True) | Rsq (All) | Rsq (VS=iSIS-MCP) | Rsq (VS=FDR) | ab (Lasso) |
|---|---|---|---|---|---|
| MSE | | | | | |
| 0 | 0 | 0.83912 | 0.00120 | 0.90069 | 0 |
| 1 | 0.00043 | 0.35468 | 0.00737 | 0.01393 | 0.00051 |
| 5 | 0.00031 | 0.01738 | 0.00210 | 0.04097 | 0.00633 |
| 10 | 0.00031 | 0.00389 | 0.00149 | 0.03771 | 0.00888 |
| 20 | 0.00033 | 0.00085 | 0.00091 | 0.03961 | 0.03112 |
| Bias | | | | | |
| 0 | 0 | 0.91602 | 0.0008 | -0.89744 | 0 |
| 1 | -0.00043 | 0.59550 | -0.04920 | -0.10082 | 0.00847 |
| 5 | 0.00006 | 0.13085 | -0.03139 | -0.19391 | -0.02649 |
| 10 | 0.00035 | 0.06012 | -0.02464 | -0.18684 | 0.01049 |
| 20 | 0.00058 | 0.02285 | -0.01445 | -0.19167 | -0.10426 |
| SD | | | | | |
| 0 | 0 | 0.00495 | 0.03467 | 0.30870 | 0 |
| 1 | 0.02068 | 0.00765 | 0.07032 | 0.06135 | 0.02090 |
| 5 | 0.01770 | 0.01595 | 0.03340 | 0.05807 | 0.07499 |
| 10 | 0.01765 | 0.01666 | 0.02965 | 0.05290 | 0.09366 |
| 20 | 0.01803 | 0.01797 | 0.02654 | 0.05362 | 0.14231 |

Table S6: The mean squared error (MSE), bias and standard deviation (SD) of $R^2_{Mediated}$ in simulation setting II (V2). Rsq (True) is $R^2_{Mediated}$ based on the true mediators $\mathbf{M}$; Rsq(All) is $R^2_{Mediated}$ based on the putative mediation variables in the initial assessment $\mathbf{M}_0$; Rsq (VS=iSIS-MCP) is $R^2_{Mediated}$ based on selected mediator set $\hat{\mathbf{M}}$ using iterative SIS; Rsq (VS=FDR) is $R^2_{Mediated}$ based on $\hat{\mathbf{M}}$ using the FDR method; ab (Lasso): product measure estimated using Lasso regression for variable selection and estimation simultaneously.

| Percentage | Rsq (True) | Rsq (All) | Rsq (VS=iSIS-MCP) | Rsq (VS=FDR) | ab (Lasso) |
|---|---|---|---|---|---|
| MSE | | | | | |
| 0 | 0 | 0.83912 | 0.00020 | 0.90069 | 0 |
| 1 | 0.00043 | 0.000371 | 0.00118 | 0.00171 | 0.03228 |
| 5 | 0.00031 | 0.000260 | 0.00097 | 0.00900 | 0.05570 |
| 10 | 0.00031 | 0.000278 | 0.00079 | 0.01491 | 0.06430 |
| 20 | 0.00033 | 0.000295 | 0.00079 | 0.02013 | 0.09642 |
| Bias | | | | | |
| 0 | 0 | 0.91602 | 0.00012 | -0.89744 | 0 |
| 1 | -0.00043 | 0.00106 | -0.02373 | -0.04087 | 0.006081 |
| 5 | 0.00006 | 0.00162 | -0.01796 | -0.09433 | -0.024859 |
| 10 | 0.00035 | 0.00051 | -0.00777 | -0.12155 | -0.017357 |
| 20 | 0.00058 | 0.00207 | 0.00415 | -0.14102 | -0.087753 |
| SD | | | | | |
| 0 | 0 | 0.00495 | 0.01424 | 0.30870 | 0 |
| 1 | 0.02067 | 0.01924 | 0.02483 | 0.00642 | 0.17955 |
| 5 | 0.01770 | 0.016034 | 0.02545 | 0.01015 | 0.23471 |
| 10 | 0.01765 | 0.01668 | 0.02695 | 0.01163 | 0.25298 |
| 20 | 0.01803 | 0.01705 | 0.02771 | 0.01561 | 0.29785 |

#### 1.6.1 Additional simulation settings

In addition to the simulating settings (V1) and (V2), we performed two other settings with $p = 15,000$ to mimic the number of genes in the real-data application:

(V3) There were $p_1 = 150$ true mediators, 1350 were $\mathbf{M}^{(2)}$ and 13,500 noise variables: $a_j \sim N(0, 0.2)$ for $j = 1, \ldots, 1500$, $b_j \sim N(0, 0.2)$ for $j = 1, \ldots, p_1$, $b_j = 0$ for $j = p_1 + 1, \ldots, 1500$, and $a_j = b_j = 0$ for $j = 1,501, \ldots, 15,000$;

(V4) There were 1500 $\mathbf{M}^{(2)}$ and 13,500 noise variables: $a_j \sim N(0, 0.2)$ for $j = 1, \ldots, 1500$, $a_j = 0$ for $j = 1501, \ldots, 15,000$, and $b_j = 0$ for $j = 1, \ldots, 15,000$.

Setting (V3) shared the same true Rsq with (V1) with 1% of the true signal. The bias of $R^2_{Mediated}$ was -0.049 and standard deviation was 0.03, while the bias of ab(lasso) was 0.06 and standard deviation was 0.13. The amount bias of $R^2_{Mediated}$ was close to (V1) with 1% of true signal (Table S5, bias=-0.049, SD=0.07) for Rsq (VS=iSIS-MCP). The bias of ab(lasso) was worse than that in (V1).

Setting (S4) shared the same true Rsq with (V1) with no true signal. The bias of $R^2_{Mediated}$ was 0.0008 and standard deviation was 0.03, while the bias of ab(lasso) 0 and standard deviation was 0. The amount bias of $R^2_{Mediated}$ was close to (V1) with 1% of true signal (Table S5, bias=-0.0008, SD=0.03) for Rsq (VS=iSIS-MCP). In the Lasso regression, none of the variables are selected, leading to the same estimation as p=1500.

### 1.7 Overall association pattern and pathway analysis in the application to the Framingham Heart Study

#### 1.7.1 Pairwise association

Fig.S3 reflects the relationship between gene expression (GE) levels and age, the marginal relationship between GE levels and the health traits, i.e., forced vital capacity (FVC) and systolic blood pressure (BP), respectively. The distribution of the coefficients of age regressed on GE skewed to the left, consistent with previous study results (De Magalhaes et al., 2009; Weindruch et al., 2002).

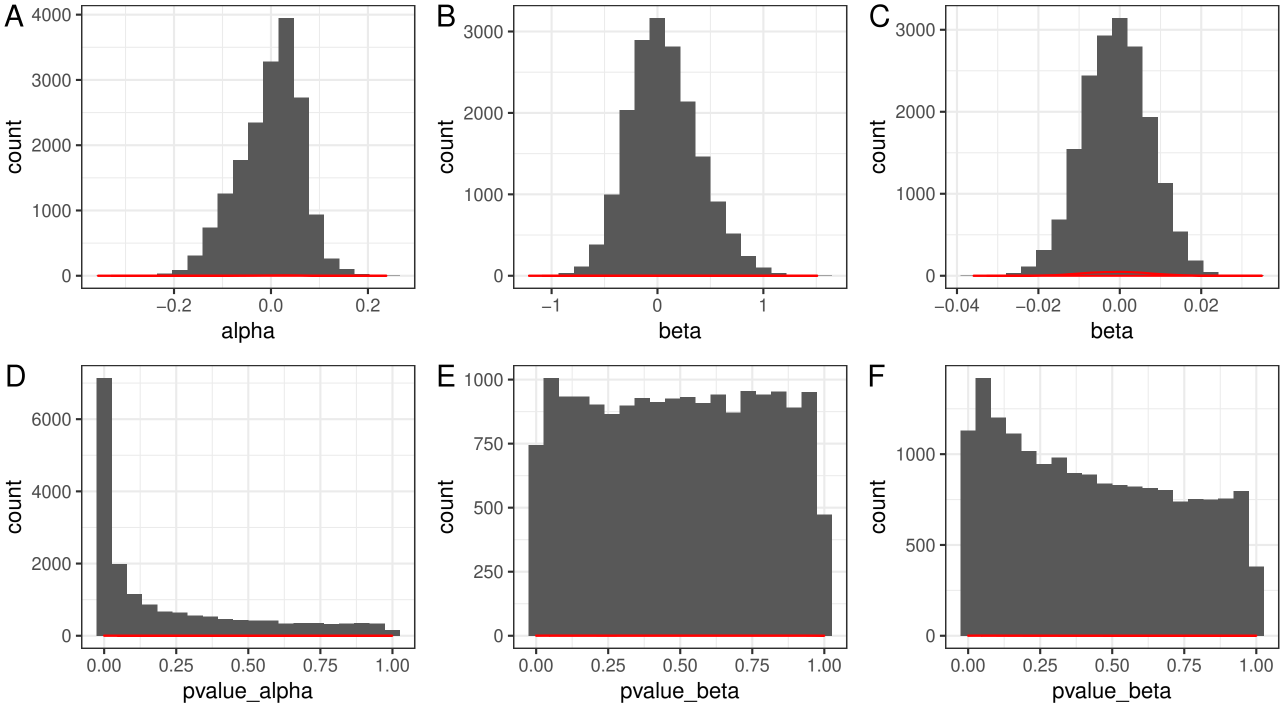


Figure S3: Coefficents of the marginal relationship in the Framingham Heart Study: A shows the distribution of the coefficients of GE levels regressed on age adjusted for covariates. B shows the distribution of the coefficients of each single GE level regressed on FVC adjusted for covariates. C demonstrates the distribution of the coefficients of systolic BP regressed on each single GE level adjusted for covariates. The corresponding p-values in plots A, B, and C are shown in plots D, E, and F, respectively.

#### 1.7.2 Pathway analysis for systolic blood pressure

For systolic BP, 182 genes whoseGEs were selected by the iterative SIS-MCP. We narrowed the gene list down to 102 genes by screening out GEs not associated with age (FDR-adjusted p-value larger than 0.1). We further performed a pathway enrichment analysis of the 102 genes that corresponded to the mediating GEs of systolic BP using the Database for Annotation, Visualization and Integrated Discovery (DAVID) v6.8 (Huang et al., 2008). One Kyoto Encyclopedia of Genes and Genomes (KEGG) pathway, Nucleotide excision repair pathway, consisting of three genes (*ERCC6*, *ERCC8*, and *CUL4B*), were statistically significant with a p-value of 0.029. The nucleotide excision repair pathway is a well-known pathway associated with age-related vascular dysfunction, which in turn is associated with hypertension (Durik et al., 2012). Furthermore, when using FDR-adjusted p-value larger than 0.2 as the cutoff point, there were 123 genes involved. Four KEGG pathways were significantly enriched at the nominal significance level of 0.05. Table S7 presents these four pathways. The first two pathways are closely related by sharing three genes. Rheumatoid arthritis is a chronic inflammatory disease and could be preceded by immune abnormalities (Goronzy et al., 2010). It could be linked to hypertension by possible sharing of pathways (Manavathongchai et al., 2013). MAPK signaling pathway was evidenced in a rat study and others to mediate in the process of aging and hypertension, related to phenotypic switching of vascular smooth muscle cells (Zhang, Zhaoxia, Ying, Jingwen, Fanxing, and Lijun, Zhang et al.; Harvey et al., 2015). It is reassuring to see the discovered pathways were supported by these previous studies.

Table S7: The top KEGG pathways for systolic blood pressure

| Name | Count | P-Value | Gene name |
|---|---|---|---|
| Collecting duct acid secretion | 3 | 0.016 | *ATP6V0A4, ATP6V0E1, ATP6V1D* |
| Rheumatoid arthritis | 4 | 0.03 | *ATP6V0A4, ATP6V0E1, ATP6V1D, FLT1* |
| MAPK signaling pathway | 6 | 0.035 | *CACNA1S, HSPA8, MAP2K5, MRAS, RAC2, MYC* |
| Nucleotide excision repair | 3 | 0.041 | *ERCC6*, *ERCC8*, *CUL4B*) |

Zhang, L., Zhaoxia, X., Ying, W., Jingwen, L., Fanxing, Z., and Lijun, S. Akt/eNOS and MAPK signaling pathways mediated the phenotypic switching of thoracic aorta vascular smooth muscle cells in aging/hypertensive rats. *Physiological research* **67,** 543–553.
